# Self-assembly of microstructured protein coatings with programmable functionality for fluorescent biosensors

**DOI:** 10.1101/2024.05.17.594773

**Authors:** Suna Jo, Erin Pearson, Donghoon Yoon, Jungkwun Kim, Won Min Park

## Abstract

Proteins, as genetically programmable functional macromolecules, hold immense potential as biocompatible self-assembling building blocks. Despite their versatility in building coating materials, it has been often hindered from programming their functionality genetically. In this study, we demonstrate a modular self-assembly of protein coatings that are genetically programmable for a biosensor application. We designed recombinant fusion protein building blocks to form microstructured coatings on diverse substrates, such as glass or polymer, through a thermally triggered liquid-liquid phase separation and an orthogonal high-affinity coiled-coil interaction. We incorporated fluorescence proteins into coatings and controlled protein density to enable fluorescence imaging and quantification in a low-resource setting. Then, we created a coating for a calcium biosensor using a genetically engineered calcium indicator protein. This protein coating served as the foundation for our smartphone-based fluorescence biosensor, which successfully measured free calcium concentrations in the millimolar range at which extracellular calcium homeostasis is maintained. Using this fluorescence biosensor, we were able to detect abnormal physiological conditions such as mild or moderate hypercalcemia. We envision that this modular and genetically programmable functional protein coating platform could be extended to the development of highly accessible, low-cost fluorescent biosensors for a variety of targets.

## 1. Introduction

Proteins have been exploited to build coating materials for a variety of applications in food packaging,^[1,2]^ drug delivery,^[3,4]^ tissue engineering,^[5,6]^ biosensors,^[7]^ biological adhesives,^[8]^ and regenerative medicine.^[3]^ Protein coatings offer a promising approach for creating functional surfaces with tailored properties such as antifouling activity^[9,10]^, selective binding affinity^[11]^, and controlled release of therapeutics.^[12,13]^ Such functional coatings include amyloid-based genetically modified protein coatings created via self-assembly, whose functionalities were programmed for biotemplating,^[14]^ biocatalysis,^[15]^ and bacterial sensors.^[7]^ Recombinant curli nanofibers were engineered to create biofilms with biocatalytic functions, where the curli nanofibers enabled modular and site-specific enzyme immobilization.^[16,17]^ In coating fabrication, microstructure formation has a significant impact on functionality and performance. Mechanical or physical properties of protein coating materials have been modulated through the control of microstructures, ^[7,18–20]^ which are influenced by the conformation, binding interaction, and topology of protein building blocks. An example is thin solid films of protein-block copolymer conjugates in which microphase-separated structures yielded high protein loading and biocatalytic activity.^[21]^ Another example includes a superhydrophobic coating with microstructures, which were manipulated by the phase transition of lysozyme. This coating presented excellent mechanical and thermal stability and the capability to drive protein crystallization.^[20]^

Recombinant proteins have been used as versatile building blocks that present excellent programmability to self-assemble biomaterials, including coating materials.^[21,22]^ Multiple genes can be recombined genetically and modularly to introduce self-assembling properties as well as a variety of functionalities, followed by protein expression and purification using well-established protocols.^[22]^ Despite their genetic programmability, processing protein materials often required through chemical modifications using organic solvents ^[23–25]^ or chemical reagents for covalent substrate attachment or cross-liking.^[21,25]^ These additional steps along with additional washing or functionalization in protein coating fabrication leave challenges in manufacturing process development. Furthermore, non-biocompatible materials and solvents can limit the functionality of recombinant proteins due to potential denaturation. Thus, a system of recombinant protein coating assembly under biocompatible conditions would be desirable for highly functional coating material development.

In this study, we demonstrate the self-assembly of modular, biocompatible protein coatings with genetic programming of functions and properties. We exploited two distinct protein motifs, an elastin-like polypeptide (ELP) and high-affinity coiled coils, to achieve coating assembly on a variety of substrate materials, manipulate microstructured morphology for high surface area, and incorporate globular functional domains to protein coatings. ELPs are tropoelastin-derived biomimetic peptides that undergo a thermally triggered liquid-liquid phase separation. The inverse phase transition results in the formation of insoluble coacervates from soluble proteins in an aqueous solution.^[26]^ Coiled coils are the α-helices that bind to each other to form a super-helical bundle. Because of their exceptional binding affinity and specificity, coiled coils have been used to program protein assemblies in various systems.^[27–29]^ We recombined ELP, coiled coils, and functional protein modules into fusion protein building blocks that form a coating material through a simple, chemical-free, and water-based process under biocompatible conditions (Figure 1).

**Figure 1.**
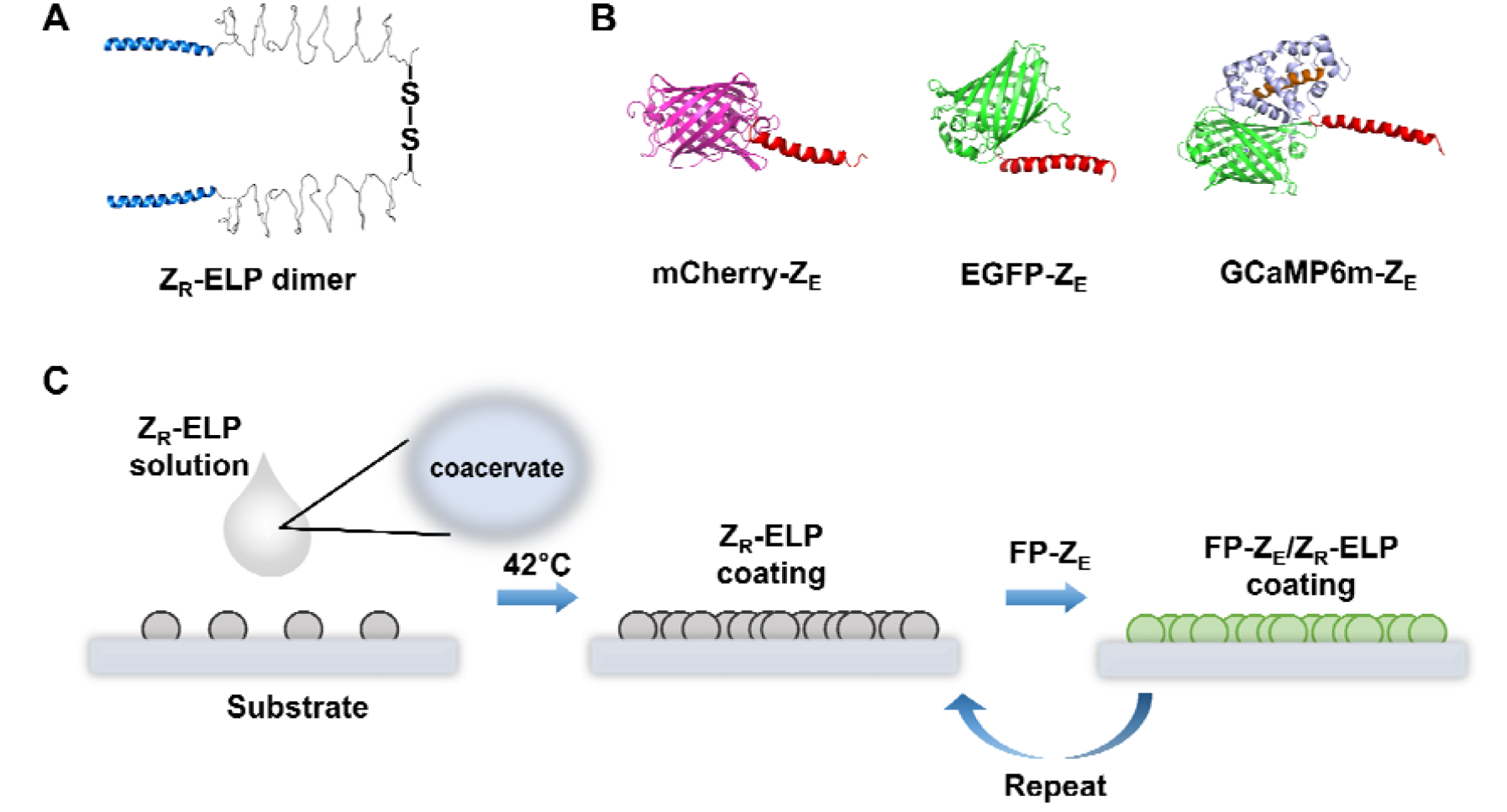
Schematic illustration of the self-assembly of a microstructured protein coating. (A) The dimeric topology of Z_R_-ELP formed through a disulfide bond. (B) Three recombinant fluorescent proteins fused with a coiled-coil tag (FP-Z_E_): mCherry-Z_E_, EGFP-Z_E_, and GCaMP6m-Z_E_. (C) Coating self-assembly process: coacervates are formed at 42°C via inverse phase transition and deposited on substrates, followed by incubation with FP-Z_E_. This process is repeated to control the density of proteins within a coating.

We used this coating platform to fabricate a smartphone-based biosensor device for the detection of hypercalcemia. We integrated a genetically encoded fluorescent calcium indicator, GCaMP, into a protein coating such that the coating exhibits fluorescence dependent upon the concentration of Ca^2+^. Employing a light-emitting diode (LED) transilluminator, and smartphone camera, we measured the fluorescence from the protein coating and quantified the Ca^2+^ concentrations in biological fluids (0.5 to 3 mM) that are relevant to mild and moderate hypercalcemia.

## 2. Results and Discussion

### 2.1 Self-assembly of a microstructured functional coating using recombinant proteins

We exploited two distinct types of protein-protein interactions to self-assemble microstructured coating materials and genetically program functionality. A pair of heterodimeric coiled-coil proteins (Z_E_:Z_R_, colon denotes heterodimerization) with a high-affinity binding interaction (dissociation constants K_D_ ∼ 10^-15^ M) ^[30,31]^ were fused to fluorescent proteins (FPs) or an ELP to construct recombinant fusion proteins, FP-Z_E_ and Z_R_-ELP.^[30]^ While an ELP triggers the self-assembly of protein-rich coacervates through a thermally responsive inverse transition, ^[30,32,33]^ the coiled coils Z_E_ and Z_R_ mediate strong binding between FP and ELP coacervate. This self-assembly system using FP-Z_E_ and Z_R_-ELP has been exploited to create protein particles in a hydrogel,^[34]^ hollow protein vesicles^[30,35,36]^, and sheet structures.^[37]^ We expressed and purified a covalently linked Z_R_-ELP dimer for protein coating self-assembly (Figures 1A and S1). A disulfide bond at the C-terminal of ELP joins the fusion protein Z_R_-ELP to one another, linking the two Z_R_ domains through an ELP_2_ linker (Z_R_-ELP_2_-Z_R_). We hypothesized that this topology is essential for the interconnection of Z_R_-ELP coacervates, which can grow into microstructured coating layers.

First, we demonstrated the self-assembly of a fluorescent coating using a fluorescent protein, mCherry. The Z_R_-ELP coacervates prepared in phosphate-buffered saline (PBS) at 25°C were first placed on a glass substrate, incubated at 42°C, and dried. The turbidity of the glass substrate measured at the wavelengths from 300 nm to 500 nm indicates the formation of a microstructured coating, which was stably adhered to the glass surface after an hour of incubation with PBS (Figure 2A). Then, the coating of Z_R_-ELP was further incubated with a solution of recombinant fusion protein mCherry-Z_E_ and washed. The fluorescence emission at 610 nm (excited at 532 nm) indicates the incorporation of mCherry-Z_E_ into the protein coating (Figure 2B), which was the affinity binding interaction between Z_E_ and Z_R_ domains. The fluorescent and confocal micrographs (Figure 2C and S2) showed the interconnection between coacervates, which resulted in the formation of microstructures adhered to glass. The fluorescence in the images (emission: 610 nm, excitation: 532 nm) further confirmed the attachment of mCherry-Z_E_ to the microstructured coating. A high-resolution acquired using field emission scanning electron microscope (FESEM) image also shows the microstructured morphology of the protein coating on glass (Figures 2D and S3). The morphology of interconnected microspheres at the size of 1.6 ± 0.5 μm was observed according to the image analysis (Figure S3). This length scale was slightly higher than the hydrodynamic diameter of mCherry-Z_E_/Z_R_-ELP coacervates measured in an aqueous solution (∼ 0.8 μm), which would be due to coalescence and growth of the coacervate microspheres on substrates. The result evidences that the microstructures evolved from the partial coalescence between the coacervate microspheres. The microstructured protein coating was stably deposited on glass without covalent conjugation or any pretreatment of the glass surfaces.

**Figure 2.**
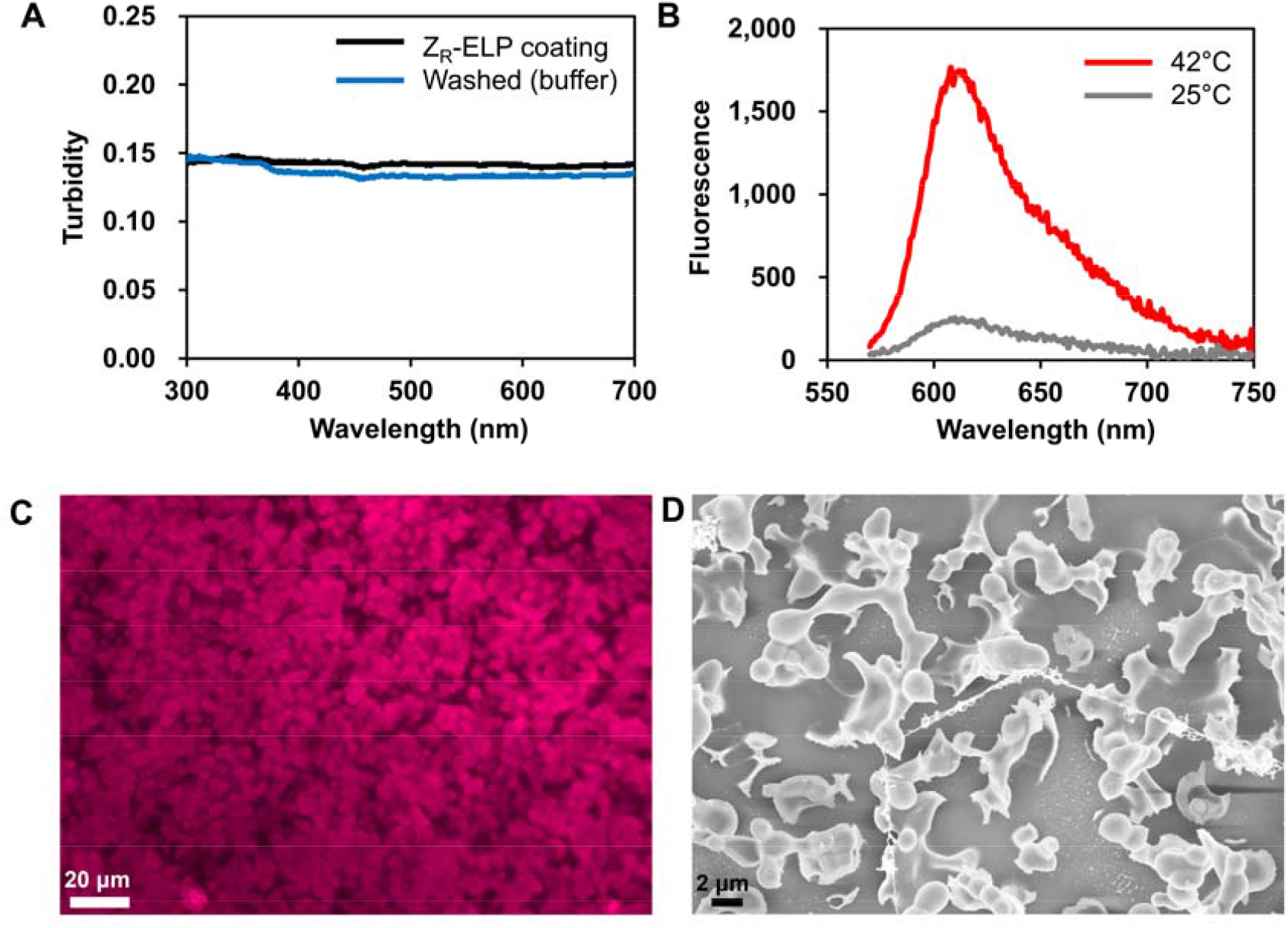
The microstructured protein coating is assembled on the glass surfaces. (A) Turbidity profile of Z_R_-ELP coating on glass (black) and incubated in buffer for 1 hr (blue). (B) Fluorescence intensities of mCherry-Z_E_/Z_R_-ELP coating prepared at 42°C (red) and 25°C (gray). (C) A fluorescence micrograph of mCherry-Z_E_/Z_R_-ELP prepared at 42°C. (D) A FESEM image of mCherry-Z_E_/Z_R_-ELP coating from panel C.

The formation of microstructures was critically influenced by temperature. In the coating sample prepared at 25°C, the formation of microstructures was not observed (Figure S2). Moreover, a remarkably lower fluorescence intensity was measured from the coating (Figure 2B). This could be due to a significant reduction in the surface area of the coating as a result of no microstructure formation. This is likely caused by structural transformation from α-helix to β-sheet within the coiled coils (Z_E_ and Z_R_). The structural transformation of coiled coils has been observed after heating, drying, or long-term storage in buffer solutions.^[38]^ Therefore, we hypothesize that the conformational transformation is facilitated near the melting temperature of Z_R_ homodimer (44°C)^[31]^, serving a critical role in stabilizing the interconnected coacervates. The Fourier Transform infrared (FT-IR) spectrum of the Z_R_-ELP coating showed reduced α-helix content and an increased fraction of β-sheet structures, indicating the transition of the secondary structure (Figure S3). The deconvoluted FT-IR spectrum in the amide I region (1600-1700 cm^-1^)^[39]^ exhibits a peak at around 1650 cm^−1^ which is indicative of the presence of 13% α-helix structures, and β-sheet structures indicated by other peaks were quantified to be 30%. In contrast, according to the circular dichroism spectra of soluble Z_R_-ELP at 25°C, the α-helix content is 35.1% and the β-strand is 13.5%, respectively.^[34]^ The results evidence the structural transformation of Z_R_ during the coating assembly.

### 2.2 Improvement in the protein coating density and stability for portable fluorescence sensing

As confirmed in Figure 2, fluorescence from the microstructured protein coating can be quantified or imaged using a spectrofluorometer or microscope. Nonetheless, a portable fluorescent biosensor application would require a simple and low-cost fluorescence reader such as a smartphone camera equipped with a light-emitting diode (LED) and filter. For this to work, the density of protein coatings on a device should be high enough to emit strong quantifiable fluorescence. To increase the protein density, we repeated the steps of coating assembly and deposition (Figure 1C). A droplet of Z_R_-ELP solution was placed and dried on a glass surface at 42°C, followed by incubation with a solution of EGFP-Z_E_.^[30]^ This process was repeated up to three times, followed by quantification of fluorescence from EGFP using a portable LED transilluminator and smartphone camera (Figure 3A). As a result, the intensity of green fluorescence after three cycles of protein deposition was increased by more than 100% than a single-deposition process. We also observed an increase in the stability of the protein coatings. The repeated deposition of proteins resulted in a reduction in protein loss after long-term incubation of a coating of Z_R_-ELP/EGFP-Z_E_ in buffer (∼ 3 hours) (Figure 3B). The coating from a single deposition showed a nearly 49% decrease in the fluorescence from the coating, while it was significantly reduced to below 10% after three or four cycles of repeated protein depositions.

**Figure 3.**
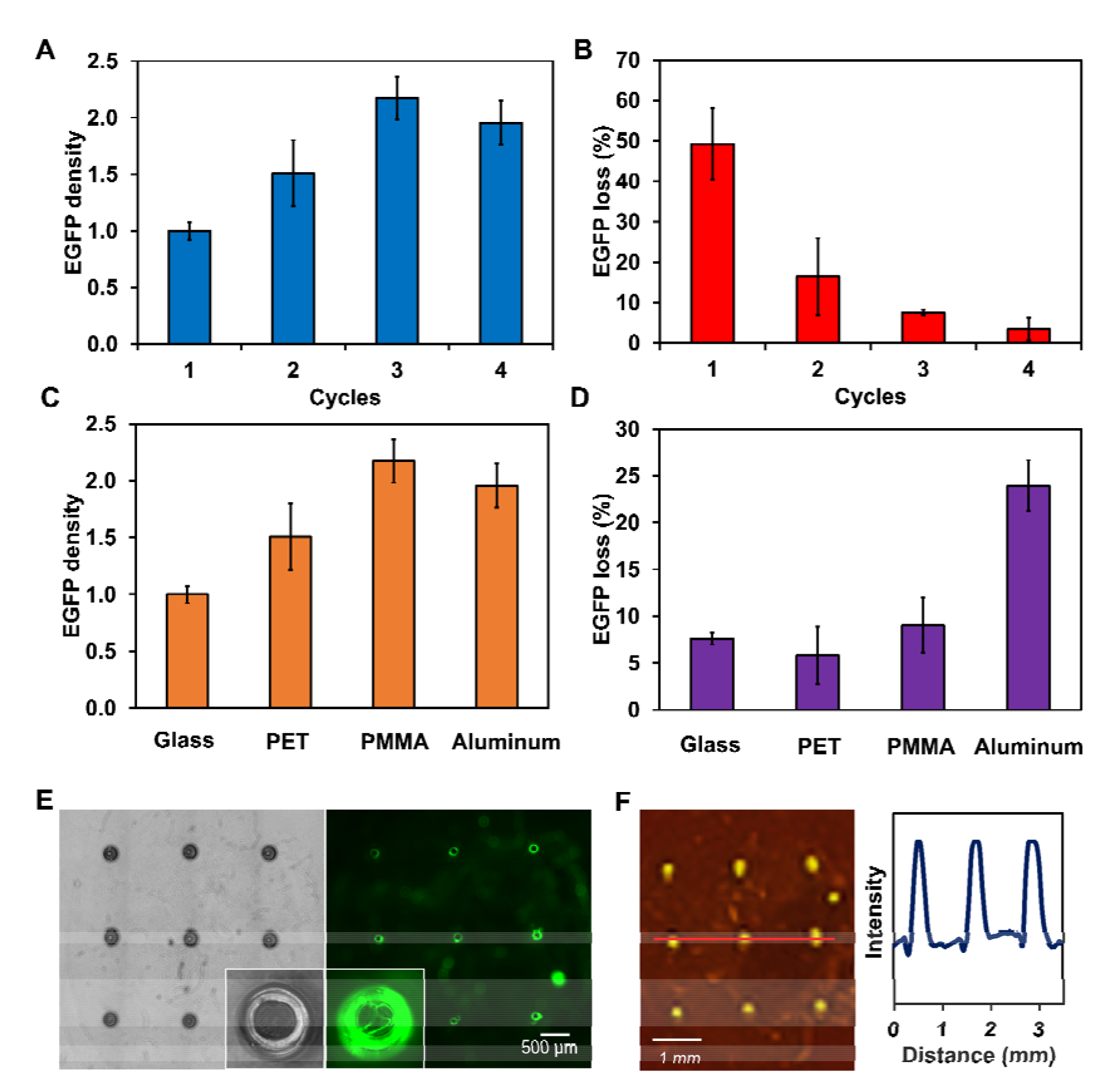
Protein density and stability of self-assembled EGFP-Z_E_/Z_R_-ELP coatings. (A) Relative protein density of EGFP in the coatings from repeated assembly and deposition cycles. The EGFP densities are normalized to the value from a single cycle. (B) Loss of EGFP from the coatings after three-hour incubation. (C) Relative protein density of EGFP on various substrates: glass, PET, PMMA, and aluminum. The EGFP densities are normalized to the value from glass. (D) Loss of EGFP from the protein coatings after a three-hour incubation. (E) Micrographs of a protein-coated micropillar array (left: bright field, right: green fluorescence) and the close-up images (insets). (F) An optical image of the protein-coated micropillar array under a transilluminator (left) and the fluorescent intensity profile of a cross-section (indicated by the red line in the left image, right).

The fluorescence intensity quantified using a smartphone camera was linearly dependent on the concentration of EGFP-Z_E_ (Figure S6). We correlated the relative fluorescence unit (RFU) and the EGFP-Z_E_ concentrations using a microplate reader, which was also correlated with the fluorescence intensity quantified from the smartphone images. The result shows a linear relationship between the fluorescence intensity of the coating, the corresponding RFU, and the protein concentration. It verifies the improvement in the density of EGFP incorporated within the coating and the stability of coating attachment.

### 2.3 Deposition of protein coatings on various substrates

Next, we conducted protein coating assembly on various substrate materials to demonstrate the versatility of our method. The coating of Z_R_-ELP/EGFP-Z_E_ from three repeated cycles of protein deposition on three different substrates other than glass, including polyethylene terephthalate (PET), polymethyl methacrylate (PMMA), and aluminum, showed higher levels of fluorescence intensity, as shown in Figure 3C. The protein coatings on the polymeric substrates (PET and PMMA) had an EGFP density of ∼150% and ∼220% relative to glass, respectively. Both were as stable as the coating on glass, according to the result from the 3-hour stability test (Figure 3D). This finding suggests that improved protein density and stability were achieved using these polymeric substrates. Although the coating on aluminum was not as stable as the one on glass, a twice higher EGFP density was achieved. Overall, the results demonstrate the versatility of our protein coating platform to be applied on various substrate materials without covalent conjugation or any pretreatment of substrates.

In biomedical applications, devices with microarchitectures are often used in combination with biological components like proteins. For example, microneedles are micron-scale medical devices used for transdermal delivery of drugs, vaccines, and other therapeutics. ^[40–42]^ To test the feasibility of our coating method on microfabricated surfaces, we conducted protein coating assembly (Z_R_-ELP/EGFP-Z_E_) on a micropillar array that is widely used for microneedles. After repeated protein coating deposition on the array of micropillars, we imaged them using a fluorescence microscope. The green fluorescence from the micropillars in the micrograph (Figure 3E) indicates the formation of the protein coating on the surface of the micropillars. We were also able to confirm that the green fluorescence could be imaged and quantified using an LED transilluminator and smartphone camera, followed by imaging analysis (Figure 3E and F). This indicates that, without functionalization, the formation of dense protein coating was achieved directly from the micropillar array. As demonstrated in Figure 3, our method can be extended to microneedles fabricated using a variety of substrates such as glass and polymers as surface treatment or functionalization is not necessary. We anticipate our approach will be a good alternative to conventional spray coating methods.^[43,44]^

### 2.4. Calcium sensing for the detection of hypercalcemia

Lastly, we demonstrated the application of our protein coating platform for a calcium biosensor using a genetically encoded calcium ion sensor protein GCaMP. The fluorescent biosensor protein is a highly effective Ca^2+^ probe, having a high signal-to-noise ratio with enhanced Ca^2+^ binding affinity (K_D_ for Ca^2+^ − 235 nM).^[45]^ GCaMP6m, an improved version of GCaMP, was recently developed by modifying the GCaMP sequence through mutagenesis to enhance its sensitivity and speed.^[46]^ The GCaMP6m protein consists of circularly-permuted GFP (cpGFP) and calmodulin (CaM), and the chicken myosin light chain kinase (M13) domains.^[47]^ Upon binding of Ca^2+^ with CaM and M13, a conformational change occurs, which alters the binding orientation of CaM and M13. This causes a change in the chemical environment surrounding the chromophore in cpGFP, which consequently increases the fluorescence intensity. Because of the critical role of intracellular Ca^2+^ in neuronal signaling, GCaMPs have been used to monitor *in vivo* neuronal activities in the mouse cortex, zebrafish, and insects, after gene transfer and transgenic expression in cells.^[46]^ Extracellular Ca^2+^ also functions as an important messenger and its increased level is associated with diseases like hypercalcemia. By assembling a coating of purified GCaMP, we used it to measure the level of Ca^2+^ in biological fluids outside cells.

We designed the recombinant fusion protein GCaMP6m-Z_E_ for the coating assembly along with Z_R_-ELP (Figure S1). After expression and purification from *Escherichia coli*, we confirmed the changes in absorbance and fluorescence intensities upon Ca^2+^ binding (Figure S7). The absorbance intensity of GCaMP6m-Z_E_ shows a significant reduction at 496 nm after the addition of a chelating agent, ethylenediaminetetraacetic acid (EDTA) at 100 µM (Figure S7A). It was immediately recovered after the addition of 100 μM of calcium chloride. Moreover, the fluorescence intensity of GCaMP6m-Z_E_ changed in response to the binding of Ca^2+^, as confirmed by the data in the presence of EDTA or CaCl_2_. (Figures S7B and S7C). The results indicate that the sensing ability of GCaMP6m was not affected by the genetic fusion of the Z_E_ coiled-coil domain.

Following the coating assembly procedure described above for Z_R_-ELP and EGFP-Z_E_, we created a protein coating that included GCaMP6m-Z_E_ on glass or PET substrates. After three cycles of coating assembly and deposition, the samples were washed and dried. The coating of Z_R_-ELP/GCaMP6m-Z_E_ on glass had a green-emitting fluorescent microstructure, as shown in the micrograph (Figure 4A). To prepare a protein coating at a Ca^2+^-free state, EDTA (100 μM) was added to a solution of GCaMP6m-Z_E_ and removed from the coating during the washing step. The protein coating was incubated with aqueous solutions at a range of Ca^2+^ concentrations, and the fluorescence from the coating was imaged using a smartphone camera, followed by image analysis to quantify the intensities. The fluorescence from the Ca^2+^-free coating was quickly recovered after exposure to Ca^2+^ in water (Figure 4C), and the relative fluorescence intensity (ΔF/F_0_) increased up to 76% when the concentrations of Ca^2+^ ranged from 1 nM to 500 µM. The apparent dissociation constant (K_D, app_) was ∼11 μM, which is shifted by two orders of magnitude from the values for soluble GCaMP6m.^[47]^ K_D, app_ can be shifted due to the higher target concentration in the micro-environment.^[48]^ The ELP-based protein coating that mimics tissue proteins is likely to provide a micro-environment, while Ca^2+^ was far above the K_D_ of soluble GCaMP6m. Therefore, this shift is most likely due to a higher Ca^2+^ concentration in the micro-environment surrounding the GCaMP probe placed within a protein-rich phase.

**Figure 4.**
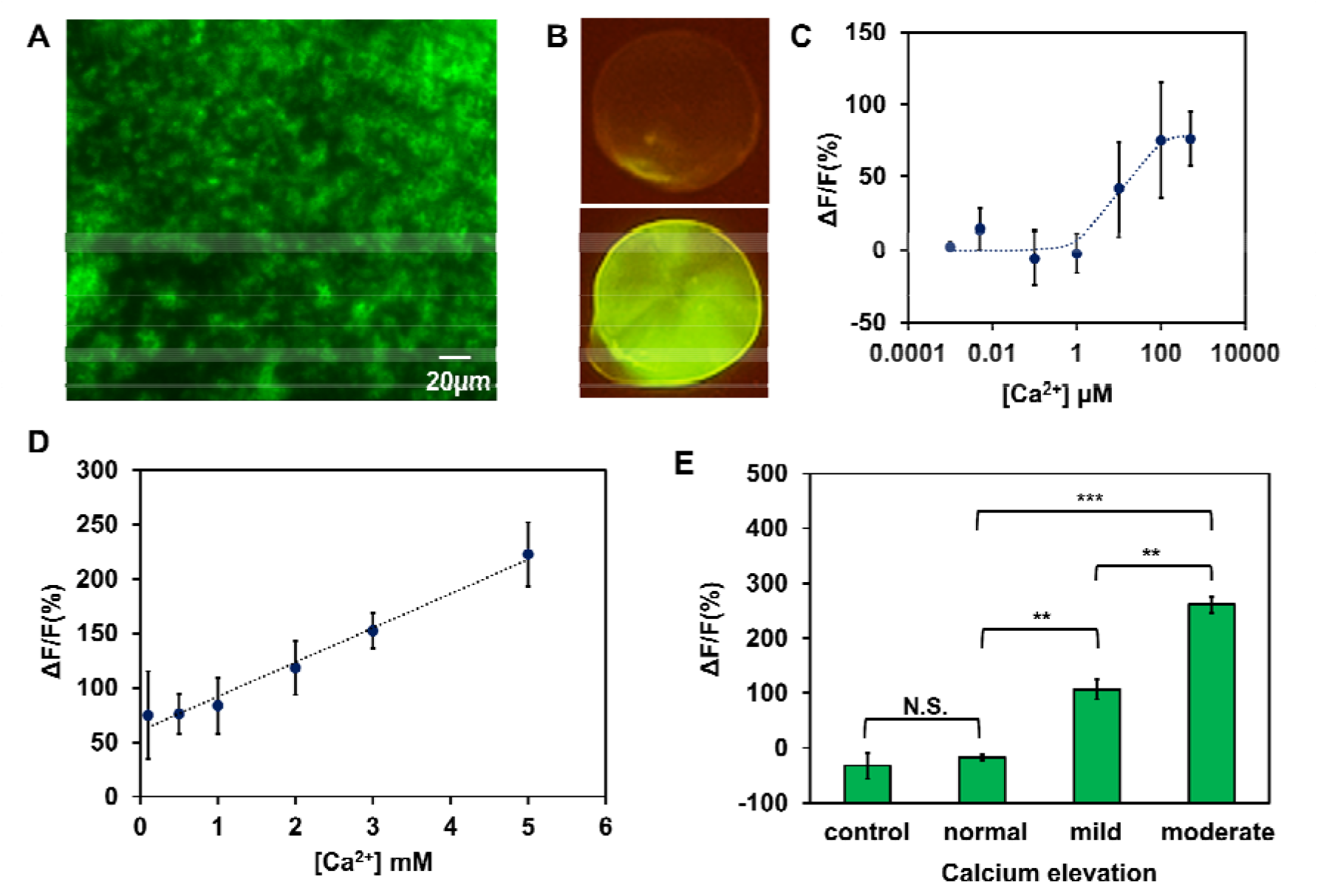
The self-assembled GCaMP6m-Z_E_/Z_R_-ELP coating. (A) A fluorescent micrograph. (B) Optical images of the coatings under a transilluminator before (top) and after incubation with 5 mM Ca^2+^ (bottom). (C) The Ca^2+^-dependent relative fluorescence intensity (ΔF/F_0_) in the range of 1 nM to 500 μM. Experimental data were fitted to a binding model (dotted line) to evaluate K_D,app._ (D) The ΔF/F_0_ curve in the range of 0.5 to 5.0 mM. Experimental data were fitted to a linear model (dotted line). (E) The ΔF/F_0_ response from Tyrode’s solution under hypercalcemia-related physiological conditions: control (no calcium), normal (1.3 mM), and mild (1.5 mM) or moderate hypercalcemia (1.8 mM). Student’s t-test: **P − 0.01, ***P − 0.001, and not significant (N.S.). The error bars in panels C, D, and E represent the standard deviation (n = 3).

The ΔF/F_0_ was also plotted against the concentrations of Ca^2+^ in the millimolar level that falls within the calcium levels in the extracellular matrix, from 0.5 to 3.0 mM (Figure 4D).^[49–51]^ At this Ca^2+^ concentration level, we observed a drastic change in the fluorescence that was linearly dependent on the concentration of Ca^2+^. The measured ΔF/F_0_ was from 76% to 223% at the Ca^2+^ concentrations of 0.5 to 5.0 mM, respectively, and an equilibrium binding model did not provide a good fit to the data. This may be due to the complex binding mechanism between GCaMP and Ca^2+^ in this concentration range and more complex binding models such as cooperativity in the ELP domain and Ca^2+^ may need to be considered after additional biophysical characterizations.^[52]^

The coating of Z_R_-ELP/GCaMP6m-Z_E_ was then used to detect hypercalcemia, which is characterized by elevated Ca^2+^ concentrations in the extracellular fluid. The normal Ca^2+^ concentration is around 1.3 mM, while mild hypercalcemia manifests with Ca^2+^ concentrations around 1.6 mM and moderate hypercalcemia exceeds 1.8 mM.^[53]^ In order to evaluate the sensing ability of our coating platform in the extracellular fluid, we conducted experiments varying Ca^2+^ concentrations in Tyrode’s solution that mimics the interstitial fluid. It is a solution with the composition of ions close to the tissue fluid and contains glucose and bovine serum albumin (BSA). The coating incubated with a solution of 1.3 mM Ca^2+^ exhibited ΔF/F_0_ of −17 ± 5 % (Figure 4E). Conversely, the coating under the mild hypercalcemia condition of 1.5 mM Ca^2+^ showed ΔF/F_0_ of 107 ± 18 %, while the ΔF/F_0_ of the sample of moderate hypercalcemia condition (1.8 mM) was 261 ± 14 %. The ΔF/F_0_ values corresponding to mild and moderate hypercalcemia were significantly higher than the normal condition (P < 0.01 for mild, P < 0.001 for moderate hypercalcemia), and the signal difference between mild and moderate hypercalcemia was also significant (P < 0.01). The reduction of fluorescence under normal or no-calcium conditions could be attributed to the interaction of Ca^2+^ with BSA.^[54]^ We confirmed from an additional experiment that the coating incubated with tris-buffered saline (TBS) with 1% BSA exhibited a reduction in fluorescence but no changes in the absence of BSA (Figure S8).

Hypercalcemia is frequently associated with several types of cancers.^[53,55]^ The regulation of calcium homeostasis is interrupted, leading to the increase of the calcium level in the bloodstream above the normal level. During the asymptotic stage, the onset of mild hypercalcemia is associated with cancer development.^[53]^ The fluorescence calcium sensor device demonstrated in this paper showed a sensing ability sufficient to detect mild or moderate hypercalcemia. The current Ca^2+^ sensors for the high concentration range (0.5–3.0 mM) are based on photonic polymers,^[56,57]^ aggregation-induced emission luminogen,^[58]^ functionalized gold nanoparticles,^[59,60]^ and a chemical fluorescence indicator,^[51]^ which require a spectrophotometer or microscope for signal measurement. In contrast, our detection method operates in a low-resource setting by using a smartphone camera equipped with an LED and light filter, which suggests a potential use for a point-of-care (POC) platform in a clinical setting. Also, our simple coating process based on protein solution casting can be extended to the manufacturing of low-cost disposable sensor platforms that were demonstrated in portable electrochemical sensors.^[61,62]^ We anticipate such a platform will facilitate a more accessible and early diagnosis of cancer in combination with the detection of other cancer-related metabolites.

## 3. Conclusion

We demonstrated a self-assembly method of protein coatings with microstructural features and programmable functionality by leveraging the modular design of recombinant proteins. The temperature-triggered coacervation through an inverse phase transition followed by non-covalent protein binding of coiled coils resulted in microstructured functional protein coatings. This simple method is applicable to various substrate materials and modular to program a functionality by incorporating globular functional protein domains like EGFP or GCaMP. We created a calcium biosensor device that quantified the Ca^2+^ level relevant to the detection of hypercalcemia conditions. This modular and versatile protein coating platform is based on a simple, chemical-free, and water-based process under biocompatible conditions. Using the genetically encoded fluorescent biosensor proteins for a variety of targets,^[63]^ we anticipate this platform will provide methods to develop a POC application with rapid and continuous monitoring of disease-associated biomarkers in a low-resource setting.

## 4. Experimental Section

### Materials

The *E. coli* strain T7 Express (BL21 derivative) was purchased from New England Biolabs. The plasmids, pQE60_Z_E_/Z_R_-ELP, pQE60_mCherry-Z_E_, and pQE60_EGFP-Z_E_, were kindly provided by Prof. Julie Champion at the Georgia Institute of Technology. Nickel nitrilotriacetic acid resin (HisPur™ Ni-NTA) was purchased from Thermo Scientific. CaCl_2_ was purchased from EMD Millipore Corporation. PBS (1X, pH 7.4) containing NaCl (137 mM), KCl (2.7 mM), NaH_2_PO_4_ (10 mM), KH_2_PO_4_ (1.8 mM), and TBS (1X, pH: 7.4) containing tris base (50 mM) and NaCl (150 mM) were prepared in the lab. Tyrode’s solution was made from lab containing NaCl (135 mM), KCl (2.8 mM), CaCl_2_ (2 mM), MgCl _2_ (1 mM), NaH_2_PO_4_ (400 μM), NaHCO_3_ (12 mM), Glucose (5.5 mM), HEPES (10 mM) and BSA. BSA was purchased from Thermo Scientific. The substrate materials of glass (0.14 mm thick), PET (0.125 mm thick), and PMMA (1.0 mm thick) were purchased from Globe Scientific Inc., Calpalmy, and Langaelex, respectively.

### Cloning

The DNA sequences encoding GCaMP6m ^[47,64]^ and the Z_E_ domain^[31]^ were recombined to create a construct encoding GCaMP6m-Z_E_. The gene fragment was synthesized, ligated into the *NcoI* and *BglII* restriction sites of the pQE60 vector (3.4kbp), and sequence-verified by GenScript. The resulting plasmid pQE60_GCaMP6m-Z_E_ was inserted into T7 Express chemically competent cells.

### Protein Expression and Purification

The recombinant fusion proteins, Z_R_-ELP, mCherry-Z_E_, and EGFP-Z_E_ were expressed and purified as previously described.^[30,34]^ The cells transformed with pQE60_GCaMP6m-Z_E_ were grown at 37□ overnight in 2X yeast extract and tryptone media supplemented with 200mg/L ampicillin. Overnight protein expression (for 18 hours) was induced at the optical density (at 600 nm) of 0.8 by adding 1.0 mM of isopropylthio-β-galactoside (IPTG). To purify GCaMP6m-Z_E_, affinity chromatography was conducted using Ni-NTA as previously described.^[30]^ After the harvest, cells were lysed through a freeze-thaw cycle and sonication, and insoluble parts were removed by centrifugation. The cleared lysate was incubated with Ni-NTA and GCaMP6m-Z_E_ was eluted by imidazole after washing. The eluted solution was dialyzed in the 1X TBS. The purity of all proteins was confirmed by sodium dodecyl sulfate-polyacrylamide gel electrophoresis.

### Protein Coating Assembly

Micropillar arrays were fabricated by UV-LED lithography, which was made using a photosensitive polymer resin.^[65]^ Substrates (Glass, PET, PMMA, and Aluminum), as well as the microfabricated micropillar arrays, were prepared on a heating block at 42 □. A 20 µl drop of Z_R_-ELP (1.7 mg/ml) was placed and dried on the substrates, followed by incubation with a solution containing 6µM of mCherry-Z_E_, EGFP-Z_E_, or GCaMP6m-Z_E_. These steps were repeated up to three times. The substrates were washed twice with 2X PBS. For a coating of GCaMP6m-Z_E_/Z_R_-ELP, a solution of GCaMP6m-Z_E_ was prepared in PBS containing 100 μM EDTA. All coatings were dried at room temperature and rinsed with TBS twice for use.

### Fluorescence Imaging and Analysis

The coatings of GCaMP6m-Z_E_/Z_R_-ELP or EGFP-Z_E_/Z_R_-ELP were placed under an LED transilluminator (BT Lab Systems) and excited at the wavelength of 470 nm. An iPhone 13 (Apple, Inc.) smartphone with a rear camera (4000x3000 pixels) was used to record fluorescence images, which were analyzed by ImageJ. Fluorescence intensity in the green channel of the images was quantified and subtracted by the intensity of background fluorescence. The quantified intensities were normalized by the background intensity and averaged by area. An equally sized area from each image was selected for the analysis.

## Acknowledgements

This research was funded by grants from the National Science Foundation (2239927) and in part by the Johnson Cancer Research Center at Kansas State University. We acknowledge Prof. Julie Champion (Georgia Institute Technology) for plasmid DNA, C. Reyes for technical assistance, and Tej Shrestha at the Nanotechnology Core in the College of Veterinary Medicine at Kansas State University for his assistance in microscope imaging.

