## Supporting Figures S1 to S7 for "Self-assembly of microstructured protein coatings with programmable functionality for fluorescent biosensors"

D. Yoon
Division of Hematology Oncology in the Department of Internal Medicine
College of Medicine, University of Arkansas for Medical Science
4301 W Markham St, Little Rock, Arkansas, 72205, USA

J. Kim
Department of Electrical Engineering, University of North Texas,
3940 N. Elm Street Ste. E255C, Denton, Texas, 76207, USA

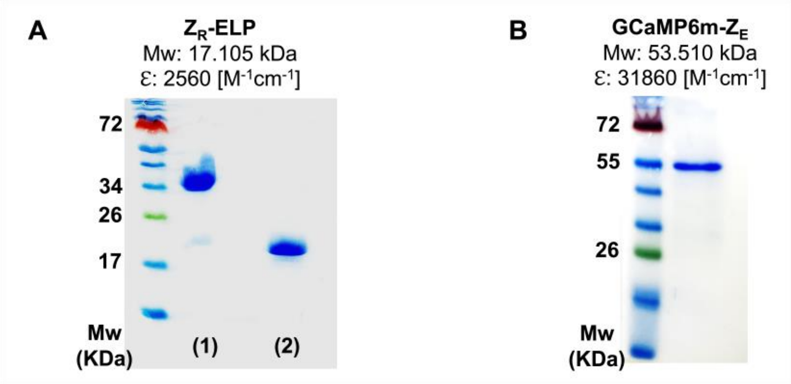


**Figure S1**. SDS-PAGE analysis of $Z_R$-ELP and GCaMP6m-$Z_E$. (A) The gel of $Z_R$-ELP samples prepared in the absence (1) or presence of the reducing agent β-mercaptoethanol (2). (B) The gel of purified GCaMP-$Z_E$. The theoretical molecular weights (Mw) and extinction coefficients (ε) are provided on top of each gel image.

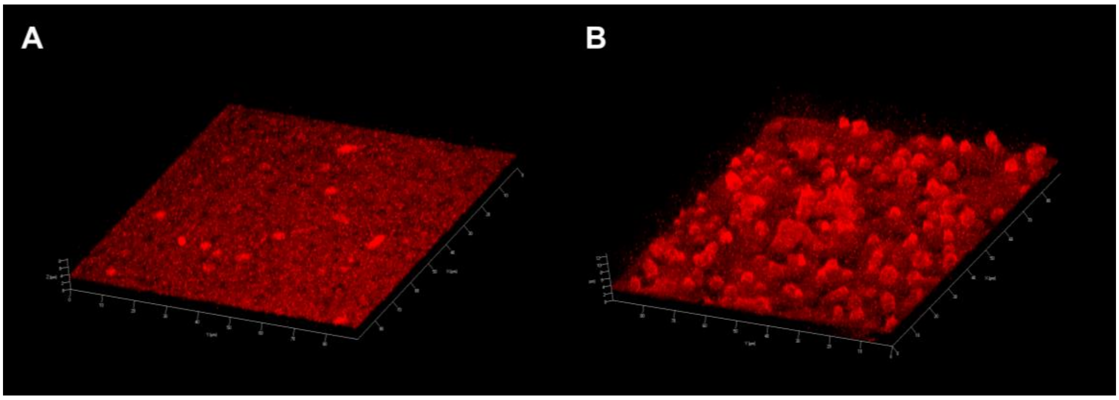


**Figure S2.** Confocal micrographs of mCherry-$Z_E$/$Z_R$-ELP coatings. Images of mCherry-$Z_E$/$Z_R$-ELP coatings on glass substrate were obtained from protein incubation at 25°C (A) and at 42°C (B).

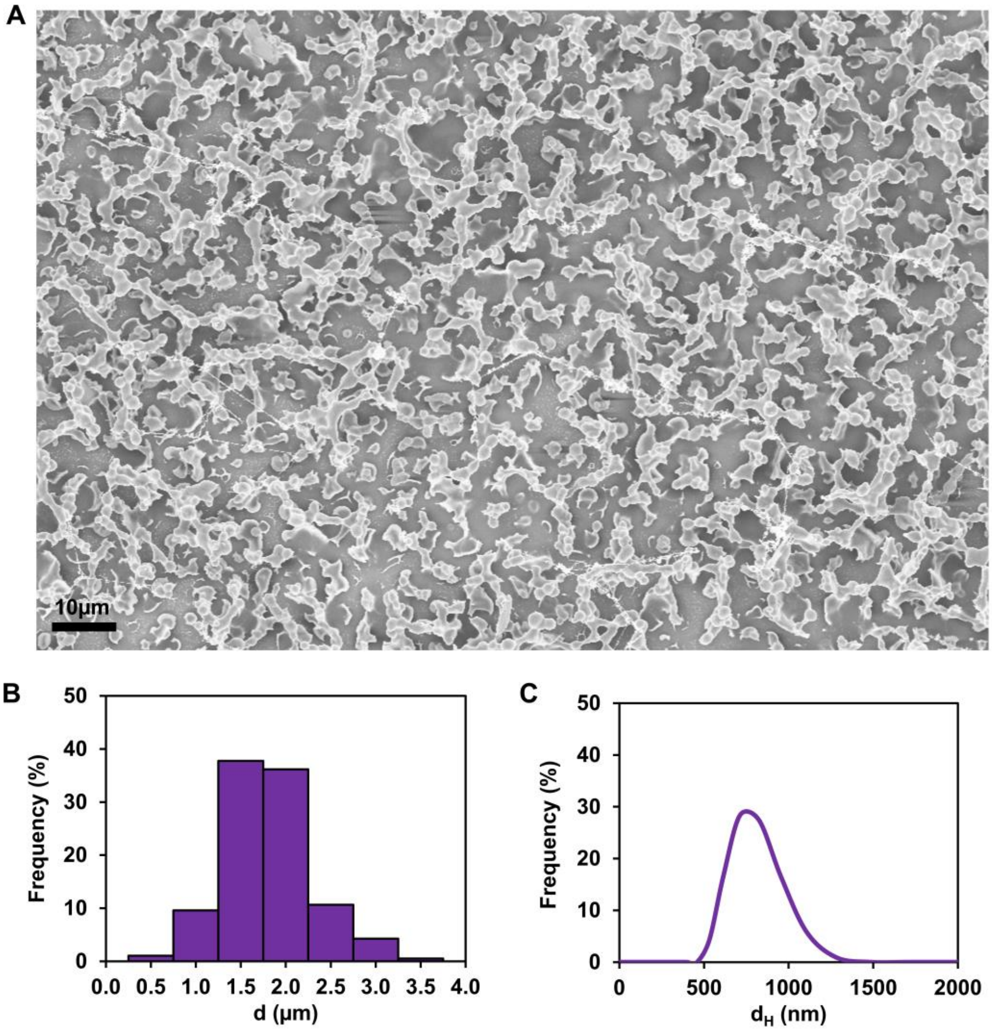


**Figure S3.** Microstructure morphology of the mCherry-$Z_E$/$Z_R$-ELP coating. (A) A FESEM image. (B) The size distribution of the coating microstructures shown in panel (A). (C) The size distribution of mCherry-$Z_E$/$Z_R$-ELP coacervates in solution measured by dynamic light scattering.

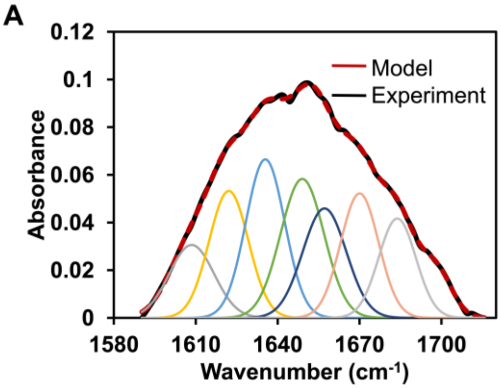


| Secondary structure | Peak position ($cm^{-1}$) | Relative area |
|---|---|---|
| β-sheet aggregates/extended strand | 1608 | 0.627 |
| β-sheet | 1622 | 0.962 |
| β-sheet | 1635 | 1.185 |
| Random coil | 1649 | 1.161 |
| α-helix | 1657 | 0.917 |
| 3(10)-helix/β-turn | 1670 | 0.910 |
| β-turn | 1683 | 0.711 |
| β-sheet/ β-turn | 1698 | 0.270 |

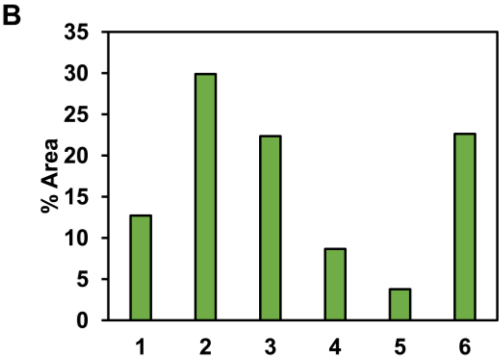


| # | Secondary structure | % area |
|---|---|---|
| 1 | α-helix | 12.7 |
| 2 | β-sheet | 29.8 |
| 3 | Random coil | 22.3 |
| 4 | β-sheet aggregates/extended strand | 8.6 |
| 5 | β-sheet/ β-turn | 3.7 |
| 6 | β-turn | 22.6 |
| | sum | 100 |

**Figure S4. (A)** Deconvolution of amide I band in FT-IR spectrum of $Z_R$-ELP coating (1600-1700 $cm^{-1}$ region). Individual components are assigned to secondary structures and relative areas are indicated in the table. (B) The secondary structure of $Z_R$-ELP coating was analyzed from the deconvolution data.

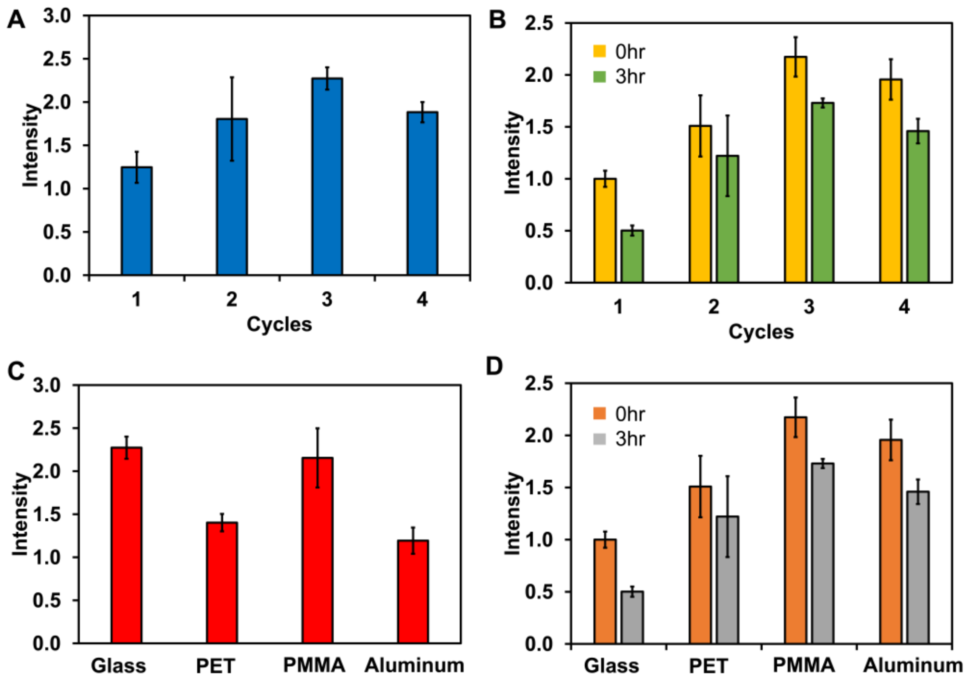


**Figure S5.** Relative fluorescence intensities from EGFP-$Z_E$/$Z_R$-ELP coatings before normalization. (A) Coatings from repeated assembly and deposition cycles. (B) Fluorescence changes from of the coatings in panel B after three-hour incubation buffer. (C) Coatings on various substrates: glass, PET, PMMA, and aluminum. (D) Fluorescence changes from of the coatings in panel C after three-hour incubation buffer.

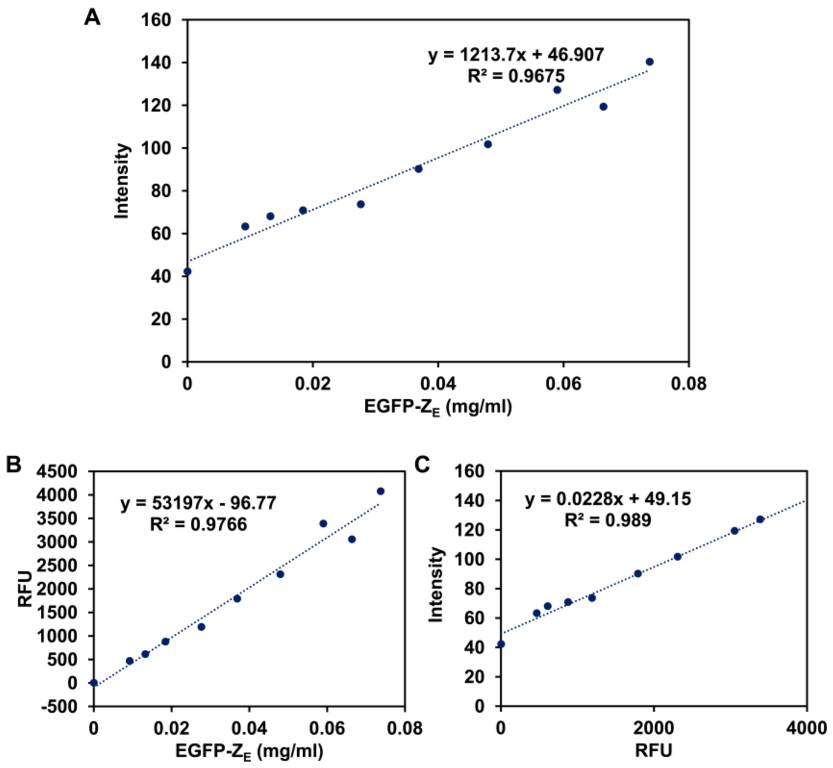


**Figure S6**. (A) Calibration curve for the fluorescence intensity from smartphone images as a function of the concentration of EGFP-$Z_E$. (B) The Relative Fluorescence Units (RFU) that correspond to the concentration of EGFP-$Z_E$. (C) The fluorescence intensity from smartphone camera images that correspond to RFU obtained from a microplate reader. The calibration curve in panel (A) was generated from the curves in panels (B) and (C).

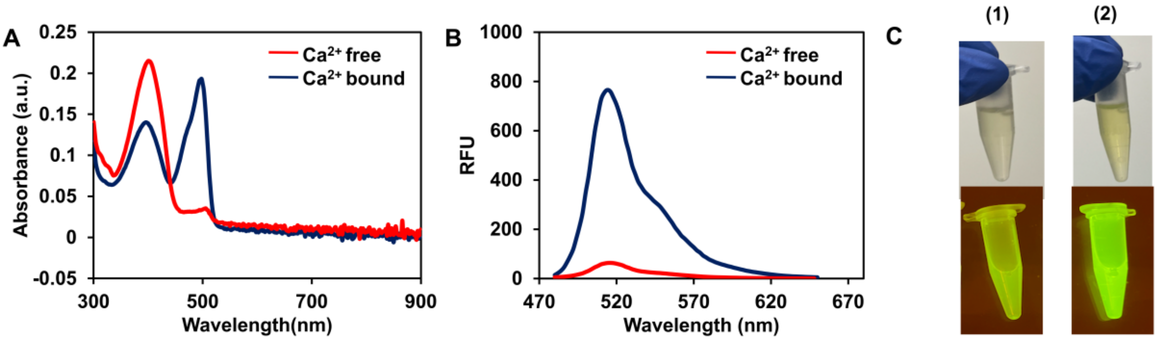


**Figure S7**. Absorbance and fluorescence changes of soluble GCaMP6m-$Z_E$ upon calcium ion binding. (A) UV-vis absorption spectra. (B) Fluorescence spectra (excitation at 470 nm). The spectra were measured in the presence of Ethylenediaminetetraacetic acid ($Ca^{2+}$-free, red) or $CaCl_2$ ($Ca^{2+}$-bound, navy blue). (C) Photographs of soluble GCaMP6m-$Z_E$ at the $Ca^{2+}$-free (1) and $Ca^{2+}$-bound state (2) under visible light (top) and transilluminator (bottom).

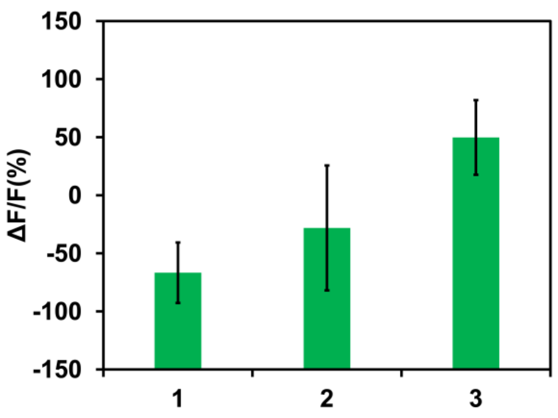


**Figure S8**. The $\Delta F/F_0$ response under hypercalcemia-related physiological conditions: (1) 1.3 mM of $CaCl_2$ in deionized water (2). 1.3 mM of $CaCl_2$ in tris buffered saline (TBS) with 1% bovine serum albumin (BSA) (3) 1.3 mM of $CaCl_2$ in TBS with no BSA.
